# Large Meta-Analysis in the CHARGE Consortium Provides Evidence For an Association of Serum Vitamin D With Pulmonary Function

**DOI:** 10.1101/144717

**Authors:** Jiayi Xu, Traci M. Bartz, Geetha Chittoor, Gudny Eiriksdottir, Ani W. Manichaikul, Fangui Sun, Natalie Terzikhan, Xia Zhou, Sarah L. Booth, Guy G. Brusselle, Ian H. de Boer, Myriam Fornage, Alexis C. Frazier-Wood, Mariaelisa Graff, Vilmundur Gudnason, Tamara B. Harris, Albert Hofman, Ruixue Hou, Denise K. Houston, David R. Jacobs, Stephen B. Kritchevsky, Jeanne Latourelle, Rozenn N. Lemaitre, Pamela L. Lutsey, George O’Connor, Elizabeth C. Oelsner, James S. Pankow, Bruce M. Psaty, Rebecca R. Rohde, Stephen S. Rich, Jerome I. Rotter, Lewis J. Smith, Bruno H. Stricker, V. Saroja Voruganti, Thomas J. Wang, M. Carola Zillikens, R. Graham Barr, Josée Dupuis, Sina A. Gharib, Lies Lahousse, Stephanie J. London, Kari E. North, Albert V. Smith, Lyn M. Steffen, Dana B. Hancock, Patricia A. Cassano

**Affiliations:** Division of Nutritional Sciences, Cornell University, Ithaca, New York, United States; Department of Biostatistics, University of Washington, Seattle, Washington, United States; Cardiovascular Health Research Unit, University of Washington, Seattle, Washington, United States; Department of Biomedical and Translational Informatics, Geisinger, Danville, Pennsylvania, United States; Icelandic Heart Association, Kopavogur, Iceland; Center for Public Health Genomics, University of Virginia School of Medicine, Charlottesville, Virginia, United States; Department of Biostatistics, Boston University School of Public Health, Boston, Massachusetts, United States; Department of Respiratory Medicine, Ghent University Hospital, Ghent, Belgium; Department of Epidemiology, Erasmus Medical Center, Rotterdam, the Netherlands; Division of Epidemiology and Community Health, University of Minnesota, Minneapolis, Minnesota, United States; Jean Mayer-U.S. Department of Agriculture Human Nutrition Research Center on Aging, Tufts University, Boston, Massachusetts, United States; Department of Respiratory Medicine, Erasmus Medical Center, Rotterdam, the Netherlands; Department of Medicine, University of Washington, Seattle, Washington, United States; Institute of Molecular Medicine, University of Texas Health Science Center at Houston, Houston, Texas, United States; Human Genetics Center, School of Public Health, University of Texas Health Science Center at Houston, Houston, Texas, United States; Children’s Nutrition Research Center, Baylor College of Medicine, Houston, Texas, United States; Gillings School of Global Public Health, Department of Epidemiology, University of North Carolina at Chapel Hill, Chapel Hill, North Carolina, United States; University of Iceland, Reykjavik, Iceland; Laboratory of Epidemiology, Demography, and Biometry, National Institute on Aging, National Institutes of Health, Department of Health and Human Services, Bethesda, Maryland, United States; Netherlands Genomics Initiative (NGI)-sponsored Netherlands Consortium for Healthy Aging, Leiden, the Netherlands; Department of Epidemiology, Harvard T.H. Chan School of Public Health, Boston, Massachusetts, United States; Department of Nutrition and Nutrition Research Institute, University of North Carolina at Chapel Hill, Kannapolis, North Carolina, United States; Sticht Center on Aging, Department of Internal Medicine, Section on Gerontology and Geriatric Medicine, Wake Forest School of Medicine, Winston-Salem, North Carolina, United States; The Pulmonary Center, Department of Medicine, Boston University, Boston, Massachusetts, United States; Department of Neurology, Boston University, Boston, Massachusetts, United States; Department of Medicine, Columbia University, New York, New York, United States; Department of Epidemiology, University of Washington, Seattle, Washington, United States; Department of Health Services, University of Washington, Seattle, Washington, United States; Kaiser Permanente Washington Health Research Institute, Seattle, Washington, United States; Institute for Translational Genomics and Population Sciences, Los Angeles Biomedical Research Institute and Department of Pediatrics at Harbor-UCLA Medical Center, Torrance, California, United States; Division of Pulmonary and Critical Care, Feinberg School of Medicine, Northwestern University, Chicago, Illinois, United States; Division of Cardiovascular Medicine, Department of Medicine, Vanderbilt University, Nashville, Tennessee, United States; Department of Internal Medicine, Erasmus Medical Center, Rotterdam, the Netherlands; Center for Lung Biology, University of Washington, Seattle, Washington, United States; Department of Bioanalysis, Ghent University, Ghent, Belgium; Division of Intramural Research, National Institute of Environmental Health Sciences, National Institutes of Health, Department of Health and Human Services, Research Triangle Park, North Carolina, United States; Behavioral and Urban Health Program, Behavioral Health and Criminal Justice Division, RTI International, Research Triangle Park, North Carolina, United States

**Keywords:** 25-hydroxyvitamin D, vitamin D, forced expiratory volume, vital capacity, respiratory function tests, smoking, human, adult, whites, African Americans

## Abstract

The role that vitamin D plays in pulmonary function remains uncertain. Epidemiological studies reported mixed findings for the association of serum 25-hydroxyvitamin D [25(OH)D] and pulmonary function. We conducted the largest cross-sectional meta-analysis of the 25(OH)D– pulmonary function association to date, based on nine European ancestry (EA) cohorts (*n*=22,838) and five African ancestry (AA) cohorts (*n*=4,290) in the CHARGE Consortium. Data were analyzed using linear models by cohort and ancestry. Effect modification by smoking status (current/former/never) was tested. Results were combined using fixed-effects meta-analysis. Mean (SD) serum 25(OH)D was 68 (29) nmol/L for EAs and 49 (21) nmol/L for AAs. For each 1 nmol/L higher 25(OH)D, forced expiratory volume in the first second (FEV_1_) was higher by 1.1 mL in EAs (95% CI: 0.9,1.3; P=2.5×10^-21^) and 1.8 mL (95% CI: 1.1,2.5; P=1.6×10^-7^) in Aas (P_race difference_=0.06), and forced vital capacity (FVC) was higher by 1.3 mL in EAs (95% CI: 1.0,1.6; P=1.1×10^-20^) and 1.5 mL (95% CI: 0.8,2.3; P=1.2×10^-4^) in AAs (P_race difference_=0.56). Among EAs, the 25(OH)D–FVC association was stronger in smokers: per 1nmol/L higher 25(OH)D, FVC was higher by 1.7 mL (95% CI: 1.1,2.3) for current smokers and 1.7 mL (95% CI: 1.2,2.1) for former smokers, compared to 0.8 mL (95% CI: 0.4,1.2) for never smokers. In summary, the 25(OH)D associations with FEV_1_ and FVC were positive in both ancestries. In EAs, a stronger association was observed for smokers compared to never smokers, which supports the importance of vitamin D in vulnerable populations.

**Cohort Funding:** This work was supported by National Institutes of Health (NIH) grant number R21 HL125574 funded by the National Heart, Lung, and Blood Institute (NHLBI) and the NIH Office of Dietary Supplements (ODS) (co-Principal Investigators [co-PIs]: DBH and PAC). The corresponding author (PAC) had full access to the data for the meta-analysis, and had final responsibility for the decision to submit for publication. No funding source had any role in the analysis of the data, the writing of the manuscript, or the decision to submit it. This work was also supported in part by R01HL077612 (PI: RGB) and by the Intramural Research Program of the National Institutes of Health (NIH), National Institute of Environmental Health Sciences (ZO1 ES043012, PI: SJL). SJL is supported by the Intramural Research Program of NIH, National Institute of Environmental Health Sciences. Infrastructure for the CHARGE Consortium is supported in part by the NHLBI grant R01HL105756.

The Age, Gene/Environment Susceptibility (AGES)–Reykjavik Study has been funded by NIH contracts N01-AG-1-2100 and 271201200022C, the National Institute on Aging (NIA) Intramural Research Program, Hjartavernd (the Icelandic Heart Association), and the Althingi (the Icelandic Parliament). The study is approved by the Icelandic National Bioethics Committee, VSN: 00-063. The researchers are indebted to the participants for their willingness to participate in the study.

The Atherosclerosis Risk in Communities Study is carried out as a collaborative study supported by NHLBI contracts HHSN268201100005C, HHSN268201100006C, HHSN268201100007C, HHSN268201100008C, HHSN268201100009C, HHSN268201100010C, HHSN268201100011C, and HHSN268201100012C. 25(OH)D measurements were conducted with the support of R01 HL103706 from the NHLBI and R01 HL103706-S1 from the NIH ODS. The authors thank the staff and participants of the ARIC study for their important contributions.

This Cardiovascular Health Study (CHS) research was supported by NHLBI contracts HHSN268201200036C, HHSN268200800007C, N01HC55222, N01HC85079, N01HC85080, N01HC85081, N01HC85082, N01HC85083, N01HC85086; and NHLBI grants U01HL080295, R01HL085251, R01HL087652, R01HL105756, R01HL103612, R01HL120393, and R01HL130114 with additional contribution from the National Institute of Neurological Disorders and Stroke (NINDS). Additional support was provided through R01AG023629 from NIA. A full list of principal CHS investigators and institutions can be found at CHS-NHLBI.org. The content is solely the responsibility of the authors and does not necessarily represent the official views of the National Institutes of Health. Vitamin D measurements were made possible by NHLBI (R01HL084443-01A2).

This work in Framingham Heart Study was supported by NHLBI’s Framingham Heart Study contract (N01-HC-25195 and HHSN268201500001I). Vitamin D measurements in the Framingham study were made possible by NIA (R01 AG14759 to SLB.).

The Health Aging and Body Composition cohort study was supported by NIA contracts N01AG62101, N01AG2103, and N01AG62106, NIA grant R01-AG028050, NINR grant R01-NR012459, and in part by the Intramural Research Program of the NIA, NIH. This research was further supported by RC1AG035835, and the serum vitamin D assays were supported by R01AG029364.

The Multi-Ethnic Study of Atherosclerosis (MESA) study is conducted and supported by NHLBI in collaboration with MESA investigators. Support for MESA is provided by contracts HHSN268201500003I, N01-HC-95159, N01-HC-95160, N01-HC-95161, N01-HC-95162, N01-HC-95163, N01-HC-95164, N01-HC-95165, N01-HC-95166, N01-HC-95167, N01-HC-95168, and N01-HC-95169 from NHLBI, UL1-TR-000040, UL1-TR-001079, and UL1-TR-001881 from NCRR, and DK063491 from the NIDDK. The MESA Lung study was supported by grants R01 HL077612, RC1 HL100543 and R01 HL093081 from NHLBI. Support for the Mineral Metabolite dataset was provided by grant HL096875.

The Rotterdam Study is funded by Erasmus Medical Center and Erasmus University, Rotterdam, the Netherlands; the Organization for the Health Research and Development (ZonMw); the Research Institute for Diseases in the Elderly (RIDE); the Dutch Ministry of Education, Culture, and Science; the Dutch Ministry for Health, Welfare, and Sports; the European Commission (DG XII), and the Municipality of Rotterdam. LL was a postdoctoral fellow of the Research Foundation—Flanders (FWO) in Brussels, Belgium. Part of this work was supported by a FWO-grant G035014N. DSM Nutritional Products AG, Kaiseraugst, Switzerland, sponsored the Vitamin D serum analyses. The authors are grateful to the study participants, the staff from the Rotterdam Study, and the participating general practitioners and pharmacists.

The Coronary Artery Risk Development in Young Adults Study (CARDIA) is supported by contracts HHSN268201300025C, HHSN268201300026C, HHSN268201300027C, HHSN268201300028C, HHSN268201300029C, and HHSN268200900041C from the National Heart, Lung, and Blood Institute (NHLBI), the Intramural Research Program of the National Institute on Aging (NIA), and an intra-agency agreement between NIA and NHLBI (AG0005).

**Author Disclosure:** Dr. Psaty serves on the DSMB of a clinical trial funded by the manufacturer (Zoll LifeCor) and on the Steering Committee of the Yale Open Data Access Project funded by Johnson & Johnson.

All other authors have no conflicts of interest. There is no commercial support or financial interest from the tobacco industry for the research presented.

The study sponsors were not involved in study design, data collection, data analysis, data interpretation, report writing, or decisions to submit the paper for publication. PAC and DBH had final responsibility for the decision to submit for publication.

**Online Supporting Material:** Supplemental table, figures, and methods are available.

**Abbreviation Footnote:**

25(OH)D: 25-Hydroxyvitamin D
AA: African Ancestry
AGES: Age, Gene, Environment, Susceptibility Study—Reykjavik, Iceland
ARIC: Atherosclerosis Risk in Communities Study
CARDIA: Coronary Artery Risk Development in Young Adults Study
CHARGE: Cohorts for Heart and Aging Research in Genomic Epidemiology Consortium
CHS: Cardiovascular Health Study
CLIA: Chemiluminescence Immunoassay
COPD: Chronic Obstructive Pulmonary Disease
EA: European Ancestry
FEV_1_: Forced Expiratory Volume in the First Second
FHS (Offspring): Framingham Heart Study—Offspring Cohort
FHS (Gen3): Framingham Heart Study—Generation 3 Cohort
FVC: Forced Vital Capacity
HABC: Health, Aging, and Body Composition Study
LC-MS/MS: Liquid Chromatography in Tandem with Mass Spectrometry
MESA: Multi-Ethnic Study of Atherosclerosis
NHANES: National Health and Nutrition Examination Survey
PFT: Pulmonary Function Test
RIA: Radioimmunoassay
RS: Rotterdam (Netherlands) Study

## INTRODUCTION

Chronic obstructive pulmonary disease (COPD), the third leading cause of mortality in the U.S.^(1)^ and among the top 10 leading causes of total years of life lost in the world^(2)^, is characterized by progressive airway obstruction. Pulmonary function tests (PFTs), as performed by spirometry, are used to quantify pulmonary function parameters including forced expiratory volume in the first second (FEV_1_) and forced vital capacity (FVC). Pulmonary function increases throughout childhood, plateaus in the 20s, and thereafter adults experience an age-related decline^(3)^. The majority of COPD cases (85%) are related to smoking^(4)^, which alters the trajectory in pulmonary function, by hindering growth, reducing peak function, and accelerating age-related decline^(5)^.

Vitamin D is proposed to have protective effects in the lungs via gene regulation^(6)^. *In vitro* studies found that 1,25-dihydroxyvitamin D, the active vitamin D metabolite, induced antimicrobial peptides for host defense in the lung and modulated airway remodeling^(7)^. In humans, 25-hydroxyvitamin D [25(OH)D] is the major vitamin D metabolite in serum, most of which forms a complex with vitamin D binding protein (∼85-90% is DBP-bound)^(8)^, and then is metabolized to 1,25-dihydroxyvitamin D [1,25-(OH)_2_D], the active steroid hormone form^(8, 9)^. Total 25(OH)D is the commonly used biomarker of vitamin D status, and it is preferred to other vitamin D metabolites, such as non-DBP-bound 25(OH)D and 1,25-(OH)_2_D, given that it is a comprehensive indicator for vitamin D stores, has a longer half-life (∼3 weeks) and is less affected by calcium^(10, 11)^. On average, African ancestry (AA) populations have lower serum 25(OH)D concentrations, due to multiple factors including genetics and skin pigmentation^(7)^, but there is evidence that AA populations have higher 1,25-(OH)_2_D levels and greater bone mineral density compared to European ancestry (EA) populations^(12)^.

Previous observational cross-sectional studies of the vitamin D–pulmonary function association in the general population reported mixed findings. Most of these studies reported a positive association between 25(OH)D and pulmonary function^(13^-^19)^, although some reported a null or inverse association^(20^-^22)^, and two others reported a positive association under certain conditions, such as only in male current smokers^(23)^ or only in overweight and obese males^(24)^. The largest previous cross-sectional study, which included two Danish cohorts (total *n* = 18,507), reported positive associations of 25(OH)D with pulmonary function^(16)^. Only one prior cross-sectional study investigated serum 25(OH)D and pulmonary function in an ancestry group other than European, and it confirmed similar positive associations in the 3,957 AA participants studied^(13)^.

The current study investigated the hypothesis that serum 25(OH)D level is positively associated with pulmonary function. We leveraged the Cohorts for Heart and Aging Research in Genomic Epidemiology (CHARGE) Consortium to include population-based data on serum 25(OH)D and pulmonary function in a harmonized analysis. Additionally, we compared the association of serum 25(OH)D and pulmonary function across EA and AA groups and investigated effect modification by cigarette smoking.

## MATERIALS AND METHODS

### Cohorts and Participants

Nine prospective cohorts in the CHARGE Consortium were included (**Table 1**). All cohorts had EA participants, and five of the cohorts had AA participants. Only one cohort [Multi-Ethnic Study of Atherosclerosis (MESA)] has participants with other ancestries, and these other ancestries were not included in this study. Among the nine cohorts, the Framingham Heart Study (FHS) had two sub-cohorts analyzed separately: the Offspring and the Third-Generation (Gen3) cohorts. Our analysis pipeline harmonized the outcome and exposure definitions, the units on all variables, and the statistical modeling. The same exclusion criteria were applied to each cohort: missing PFTs, unacceptable PFTs using the American Thoracic Society and European Respiratory Society criteria for acceptability, missing serum 25(OH)D, serum 25(OH)D > 374.4 nmol/L (or 150ng/mL, leading to removal of a single outlier)^(25)^, or missing on other covariates (**Supplemental Table 1**).

**Table 1.**
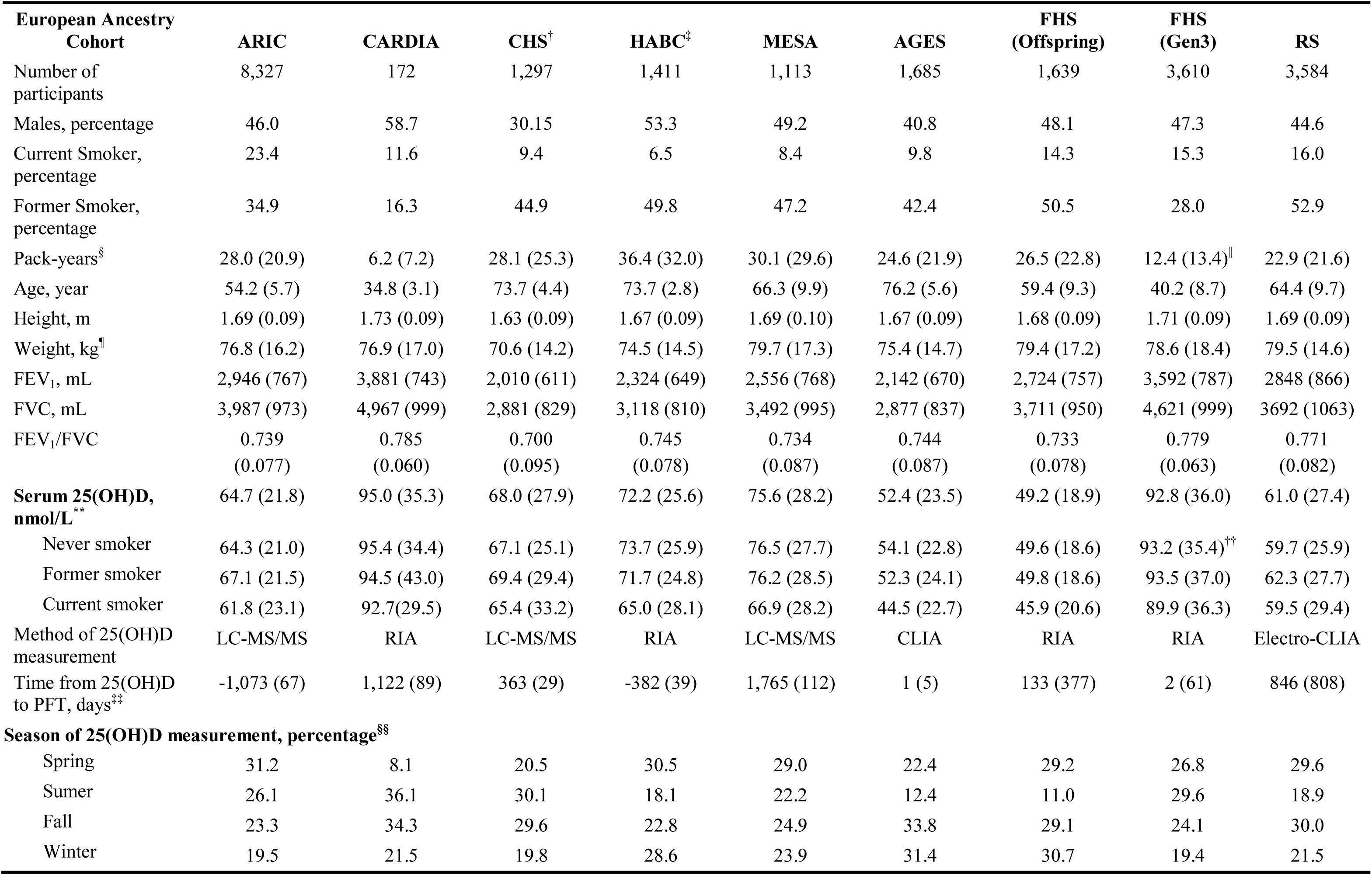
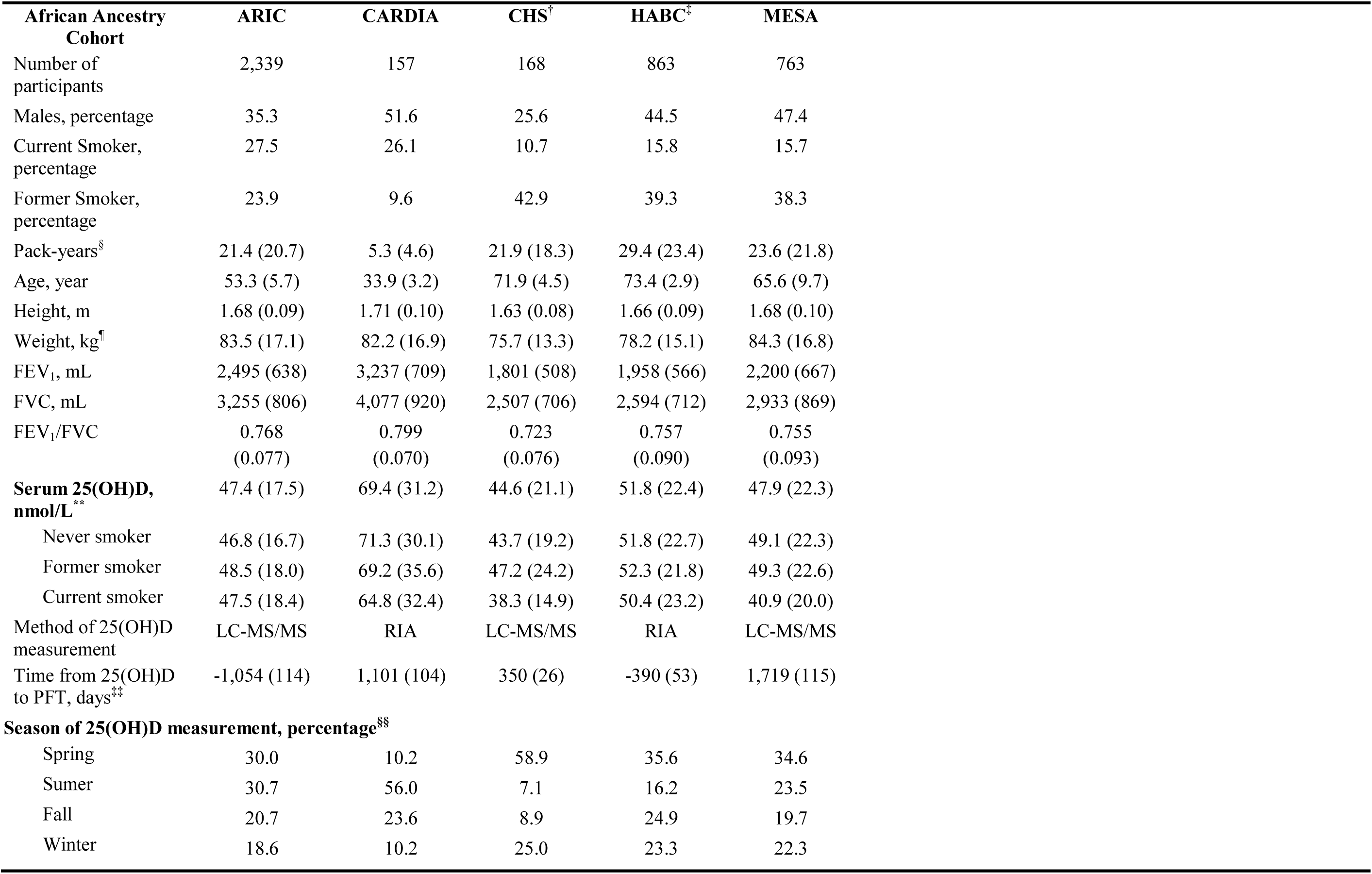
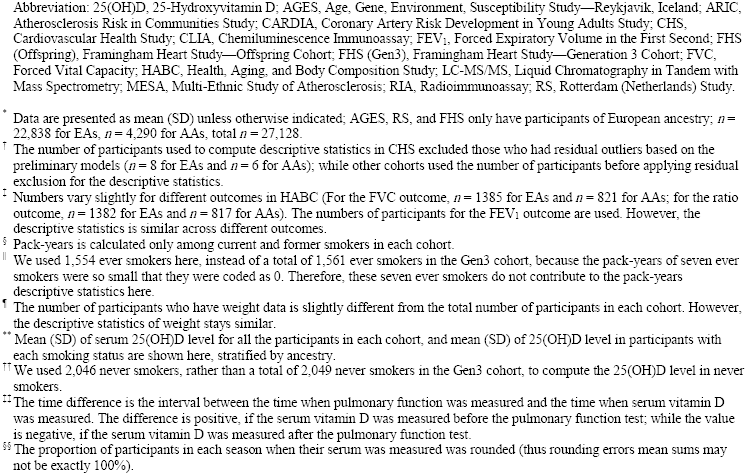
Cross-sectional participant characteristics of each cohort in the CHARGE Consortium (*n* = 27,128)*

### Outcome and Exposure Assessment

Pre-bronchodilator pulmonary function outcomes (FEV_1_, FVC, and FEV_1_/FVC), which have similar accuracy as post-bronchodilator measures for long-term outcomes^(26)^, were measured in each cohort using standardized methods defined by the American Thoracic Society/European Respiratory Society criteria (**Supplemental Table 2**). The methods used to measure 25(OH)D varied by cohort (**Supplemental Table 2**). Three cohorts, including MESA, the Atherosclerosis Risk In Communities (ARIC) study, and the Cardiovascular Health Study (CHS), used the current reference method, liquid chromatography in tandem with mass spectrometry (LC-MS/MS); three cohorts, including FHS, the Coronary Artery Risk Development in Young Adults (CARDIA) study, and the Health, Aging, and Body Composition (HABC) study, used radioimmunoassay (RIA); one cohort, the Age, Gene, Environment, Susceptibility Study— Reykjavik, Iceland (AGES), used chemiluminescence immunoassay (CLIA); and one cohort [the Rotterdam Study (RS)] used electro-CLIA. Only MESA calibrated the serum 25(OH)D measurement against the standard reference material 972^(27)^, which reflects the calendar time of the measurements in the cohorts, most of which occurred before the availability of the standard reference material (**Supplemental Table 3**). CARDIA measured serum vitamin D in a subset of participants included in an ancillary study of bone mineral homeostasis^(28)^. For the remaining cohorts, measurements of the outcome and exposure variables were planned for either the full cohort or a random sample (**Supplemental Table 1**). Continuous variables were used for serum 25(OH)D and pulmonary function to capture the association of 25(OH)D on PFTs across the broad distribution of ranges in the cohorts.

As shown in **Table 1**, among nine cohorts, four [AGES, CHS, FHS-Offspring, and FHS-Gen3] had a mean time difference of less than one year in the PFT measurements and the preceding 25(OH)D measurement, and the greatest mean time difference between 25(OH)D and PFT measurement was < 5 years [MESA]. Participants in ARIC and HABC had blood drawn for serum 25(OH)D after their PFT measure, but within 3 years.

Other covariates, including smoking status, pack-years (number of packs of cigarettes smoked per day times the number of years smoked), height, weight, and age, were measured concurrently with pulmonary function, except for CHS, which assessed covariates concurrent with the serum 25(OH)D measure, but within 1 year of the PFT measurement (**Supplemental Table 3**). All data collection and analysis was approved by the Institutional Review Board at each cohort’s respective institution. Spirometry measures are available on the database of Genotypes and Phenotypes via accession numbers as follows: ARIC (phs000280), CARDIA (phs000285), CHS (phs000287), FHS (phs000007), and MESA (phs000209). Serum vitamin D measures are also available at the same accession numbers for CHS, FHS, and MESA.

### Statistical Analysis in Individual Cohorts

All analyses were first conducted independently in each cohort, stratified by ancestry, given the lower mean serum 25(OH)D level in AA participants^(7)^. For FEV_1_ and FEV_1_/FVC, models were adjusted for smoking status, pack-years, height, height squared, age, age squared, sex, season of blood draw, and study center (if applicable); for FVC, the model was further adjusted for weight. Residual outliers, identified using the studentized residuals of the linear models (**Supplemental Methods** for more details), were excluded from all models. The model was extended to test the interaction between 25(OH)D and smoking status [never (reference group), former, and current smokers].

### Meta-Analysis

Fixed-effects meta-analysis was conducted for the association of serum 25(OH)D on each PFT outcome for each ancestry group, using inverse variance weighting, with heterogeneity assessed via the I^2^ statistic^(29)^. The comparison of meta-analyzed coefficients of the 25(OH)D–PFT associations for the two ancestry groups was conducted using a Z test^(30)^. Meta-analysis of the interaction terms of 25(OH)D with smoking status was also performed (**Supplemental Methods** for more details).

Meta-regression was conducted to explore the potential causes of heterogeneity in the primary meta-analysis of serum 25(OH)D on FEV_1_ (or FVC) in EAs. Modifiers were tested individually in the meta-regression models to investigate heterogeneity; modifiers included factors that could vary between cohorts, such as proportion of ever, current, and former smokers, mean 25(OH)D level, assay method for serum 25(OH)D, time between 25(OH)D and PFT measures, and mean age of participants in each cohort. The two-sided type I error was examined at 0.05 for all analyses. Meta-analysis and meta-regression were conducted using the metafor package (version 1.9-8) in R (version 3.2.3., R Foundation for Statistical Computing, Vienna, Austria).

## RESULTS

We studied 22,838 EA and 4,290 AA participants. EA participants had higher FEV_1_, FVC, and serum 25(OH)D than AA participants in each cohort, while FEV_1_/FVC was similar across ancestry groups (**Table 1** and **Supplemental Figure 1**). CARDIA and FHS-Gen3 were younger than the seven other cohorts, with consequently lower pack-years smoked in ever smokers. Across all cohorts, among EA participants, 17% were current smokers and 40% were former smokers; among AA participants, 22% were current smokers and 30% were former smokers. The serum 25(OH)D level was highest among never smokers [mean(SD) = 70(30) nmol/L], followed by former smokers [67 (29) nmol/L], and current smokers [64 (29) nmol/L] in EAs, while the trend was less obvious in AAs [49 (21) nmol/L in current smokers, 50 (21) nmol/L in former smokers, and 48 (21) nmol/L in never smokers]. The mean (SD) of serum 25(OH)D for EA participants across nine cohorts was 68 (29) nmol/L and for AA participants across five cohorts the mean (SD) was 49 (21) nmol/L.

**Figure 1.**
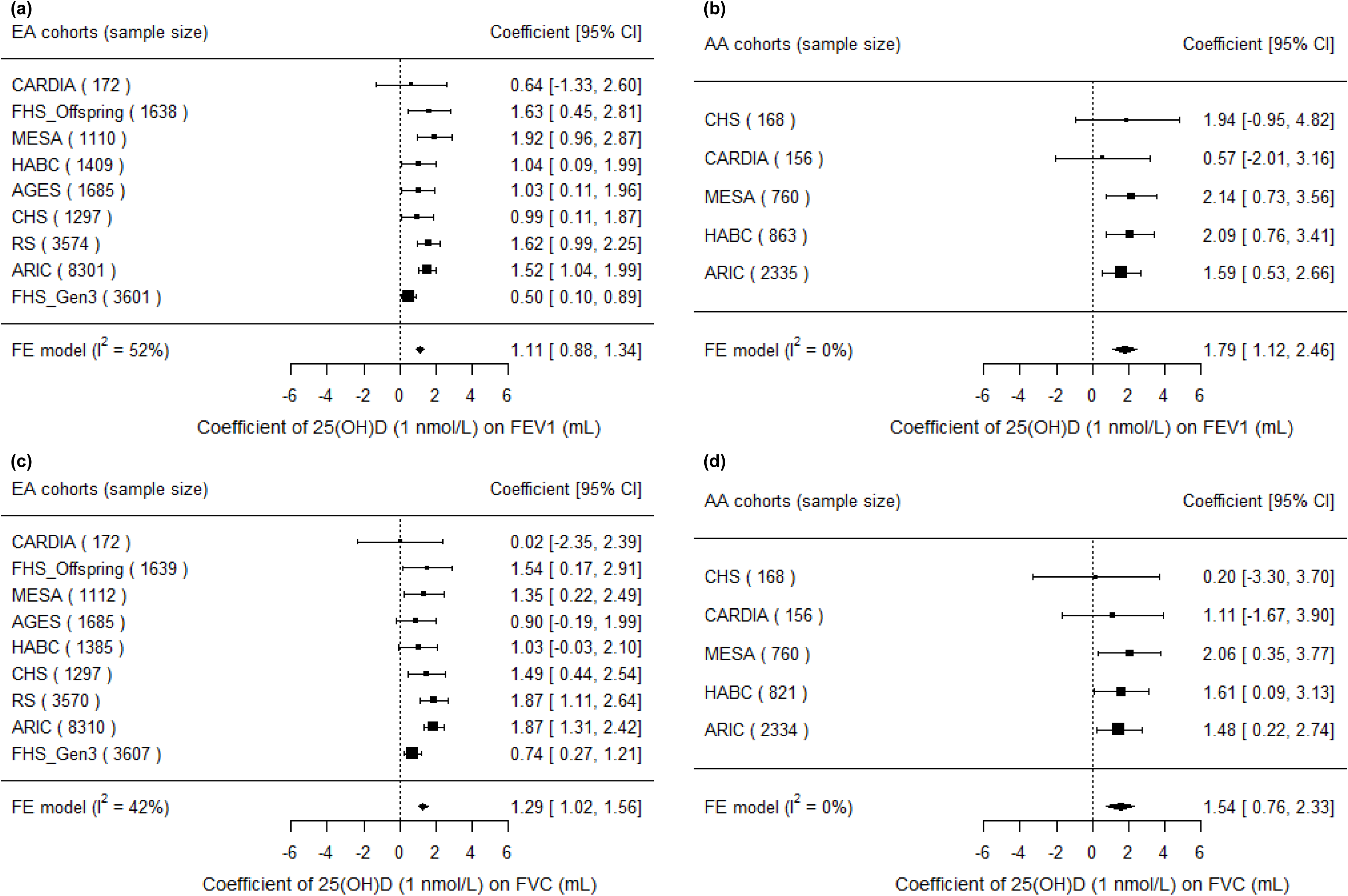
Forest plots of the meta-analysis of serum 25(OH)D on FEV_1_ and FVC across cohorts in the CHARGE Consortium, stratified by participant ancestry. Associations are presented for serum 25(OH)D on (A) FEV_1_ in European ancestry cohorts (*n* = 22,787). (B) FEV_1_ in African ancestry cohorts (*n* = 4,282). (C) FVC in European ancestry cohorts (*n* = 22,777). (D) FVC in African ancestry cohorts (*n* = 4,239). β (unit: mL) denotes the coefficient from the fixed-effects meta-analysis for serum 25(OH)D on the pulmonary function outcome per 1 nmol/L increment of 25(OH)D, with its 95% confidence interval. Cohorts findings were ordered from the least to the most precise, and heterogeneity is presented (I^2^). Abbreviation: 25(OH)D, 25-Hydroxyvitamin D; AA, African Ancestry; AGES, Age, Gene, Environment, Susceptibility Study—Reykjavik, Iceland; ARIC, Atherosclerosis Risk in Communities Study; CARDIA, Coronary Artery Risk Development in Young Adults Study; CHS, Cardiovascular Health Study; CI, Confidence Interval; EA, European Ancestry; FE, Fixed-Effects; FEV_1_, Forced Expiratory Volume in the First Second; FHS (Offspring), Framingham Heart Study—Offspring Cohort; FHS (Gen3), Framingham Heart Study—Generation 3 Cohort; FVC, Forced Vital Capacity; HABC, Health, Aging, and Body Composition Study; MESA, Multi-Ethnic Study of Atherosclerosis; RS, Rotterdam (Netherlands) Study.

Regression coefficients (β) and standard errors (SE) calculated within each cohort per 1 nmol/L 25(OH)D are presented in the figures. Additionally, to put the magnitude of the 25(OH)D–PFT associations in terms relevant to public health, the meta-analyzed regression coefficients were multiplied by 10 nmol/L 25(OH)D, which is about half of the standard deviation (SD) of the 25(OH)D distribution.

Meta-analysis **(Figure 1**) revealed a consistently positive association of serum 25(OH)D with the PFT outcomes, FEV_1_ and FVC, in both ancestry groups. To put these findings into context, a 10 nmol/L (∼0.5 SD) higher 25(OH)D was associated with 11.1 mL higher FEV_1_ in EAs (P = 2.5×10^-21^) and 17.9 mL higher FEV_1_ in AAs (P = 1.6×10^-7^). Similarly, for a 10 nmol/L higher 25(OH)D, FVC was higher by 12.9 mL in EAs (P = 1.1×10^-20^) and by 15.4 mL in AAs (P = 1.2×10^-4^). The magnitudes of the 25(OH)D–PFT associations did not differ significantly between the two ancestry groups (P = 0.06 and P = 0.56 for FEV_1_ and FVC, respectively). The association of serum 25(OH)D with FEV_1_/FVC reached statistical significance only in EAs (P = 0.0013), and the magnitude was negligible; a 10 nmol/L higher 25(OH)D was associated with a ratio being lower by 0.0055% (**Supplemental Table 4 and Supplemental Figure 2** for ancestry- and cohort-specific findings).

**Figure 2.**
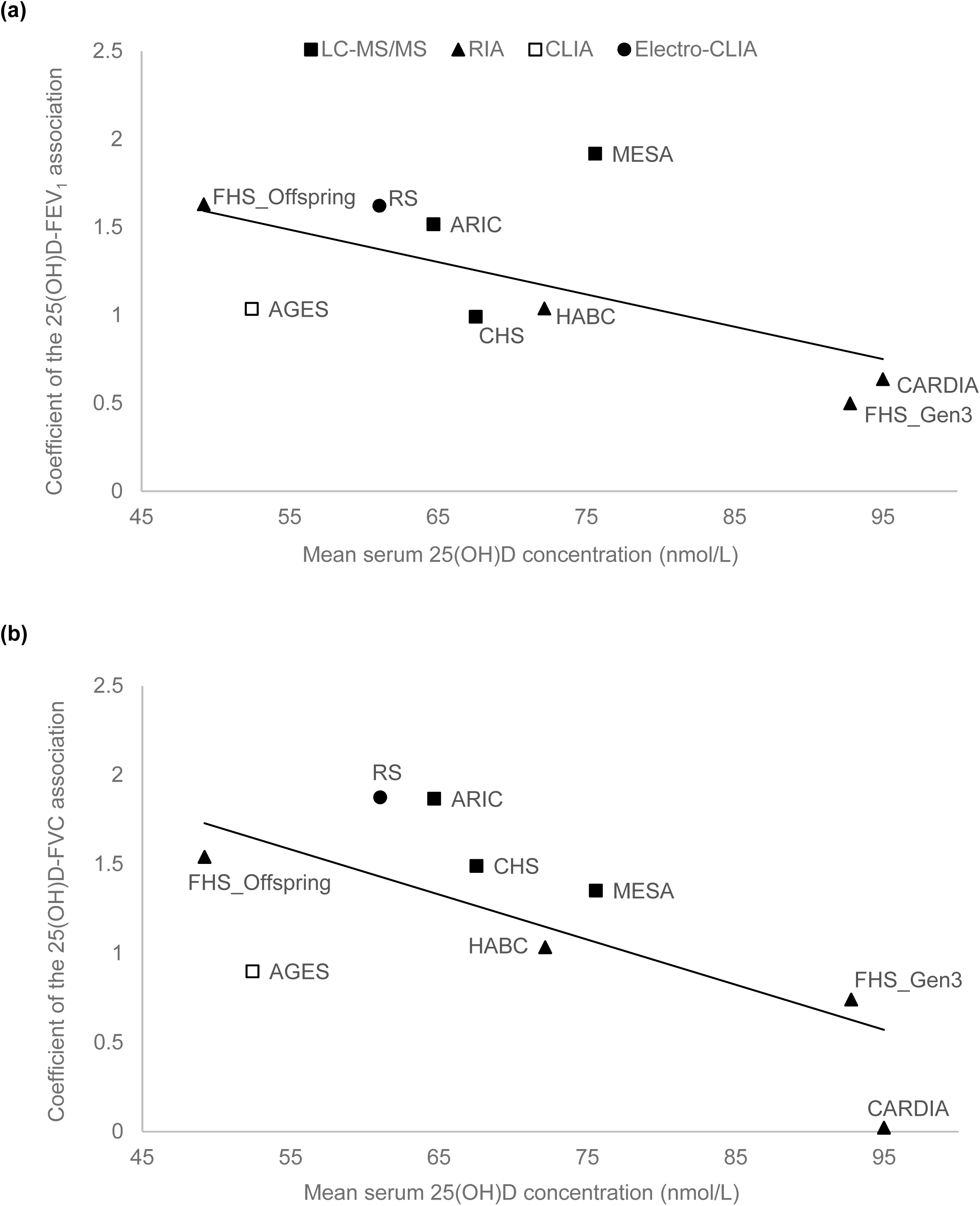
Meta-regression of mean serum 25(OH)D levels against the association estimates of 25(OH)D with PFT in nine European ancestry cohorts in the CHARGE Consortium. (A) FEV_1_ outcome (coefficient unit: mL per 1 nmol/L 25(OH)D), and (B) FVC outcome (coefficient unit: mL per 1nmol/L 25(OH)D). The modifier is mean serum 25(OH)D level of each nine cohorts. A linear regression line is present for each sub-figure, with a meta-regression p-value of 0.0006 for the FEV_1_ outcome, and 0.005 for the FVC outcome. The figure also shows the measurement method for the serum 25(OH)D assay (legend shows symbols for each of the 4 assay methods). Abbreviations: 25(OH)D, 25-Hydroxyvitamin D; AGES, Age, Gene, Environment, Susceptibility Study—Reykjavik, Iceland; ARIC, Atherosclerosis Risk in Communities Study; CARDIA, Coronary Artery Risk Development in Young Adults Study; CHS, Cardiovascular Health Study; CLIA, Chemiluminescence Immunoassay; FEV_1_, Forced Expiratory Volume in the First Second; FHS (Offspring), Framingham Heart Study—Offspring Cohort; FHS (Gen3), Framingham Heart Study—Generation 3 Cohort; FVC, Forced Vital Capacity; HABC, Health, Aging, and Body Composition Study; LC-MS/MS, Liquid Chromatography in Tandem with Mass Spectrometry; MESA, Multi-Ethnic Study of Atherosclerosis; PFT, Pulmonary Function Test; RIA, Radioimmunoassay; RS, Rotterdam (Netherlands) Study.

In the main-effect meta-analysis of serum 25(OH)D on pulmonary function, there was low to moderate heterogeneity in the EA cohorts, and low heterogeneity in the AA cohorts (**Figure 1, Supplemental Figure 2**). We used meta-regression to explore potential causes of moderate heterogeneity in the meta-analysis of 25(OH)D on FEV_1_ and FVC in the EA cohorts. Cohorts with lower mean 25(OH)D concentration had stronger 25(OH)D–PFT associations (**Figure 2**). The proportion of ever smokers and of former smokers had significant linear associations with the 25(OH)D–PFT coefficients (**Supplemental Figure 3**), and these two variables were both highly correlated with mean 25(OH)D levels (Pearson’s r > 0.75 for all pairwise correlations). The 25(OH)D–PFT association in EA cohorts varied by 25(OH)D assay method (meta-regression p < 0.02); the association was attenuated in cohorts using RIA compared to cohorts using LC-MS/MS (pairwise p < 0.005, **Supplemental Figure 4**). Mean age of each cohort was a significant positive modifier of the 25(OH)D–FEV_1_ association, while time difference between 25(OH)D and spirometry measures did not affect the 25(OH)D–PFT association (**Supplemental Figure 3**).

**Figure 3.**
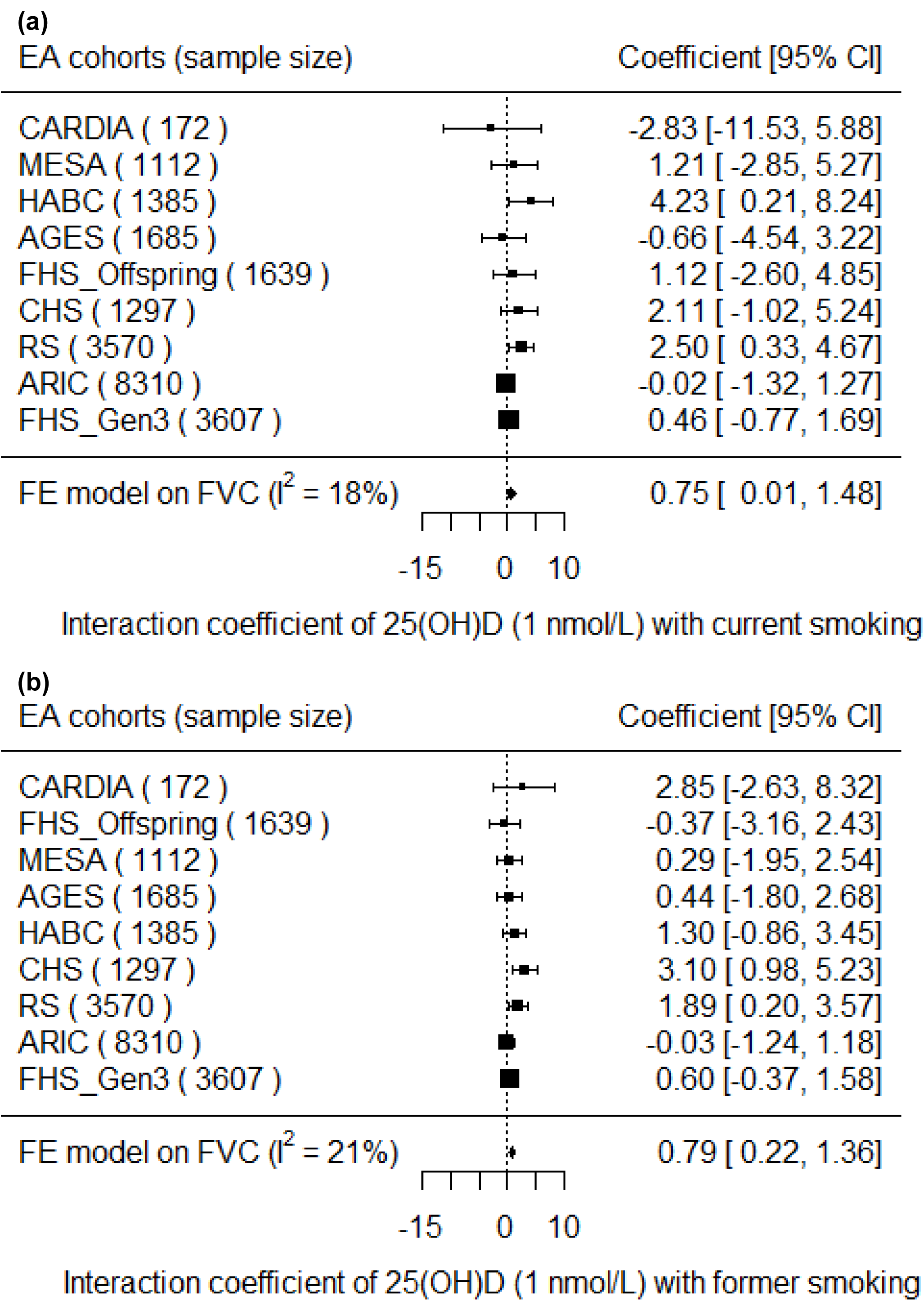
Forest plots of the interaction meta-analysis of serum 25(OH)D and smoking status on FVC in the European ancestry cohorts in the CHARGE Consortium (*n* = 22,777). (A) Current Smokers and (B) Former Smokers. β (unit: mL) denotes the interaction term coefficient of 25(OH)D and smoking status on FVC from the fixed effects meta-analysis, per 1 nmol/L increment of 25(OH)D, with its 95% confidence interval. Cohorts were ordered from the least to the most precise, and heterogeneity is presented (I^2^). Abbreviation: 25(OH)D, 25-Hydroxyvitamin D; AGES, Age, Gene, Environment, Susceptibility Study—Reykjavik, Iceland; ARIC, Atherosclerosis Risk in Communities Study; CARDIA, Coronary Artery Risk Development in Young Adults Study; CHS, Cardiovascular Health Study; CI, Confidence Interval; EA, European Ancestry; FE, Fixed-Effects; FHS (Offspring), Framingham Heart Study—Offspring Cohort; FHS (Gen3), Framingham Heart Study— Generation 3 Cohort; FVC, Forced Vital Capacity; HABC, Health, Aging, and Body Composition Study; MESA, Multi-Ethnic Study of Atherosclerosis; RS, Rotterdam (Netherlands) Study.

**Figure 4.**
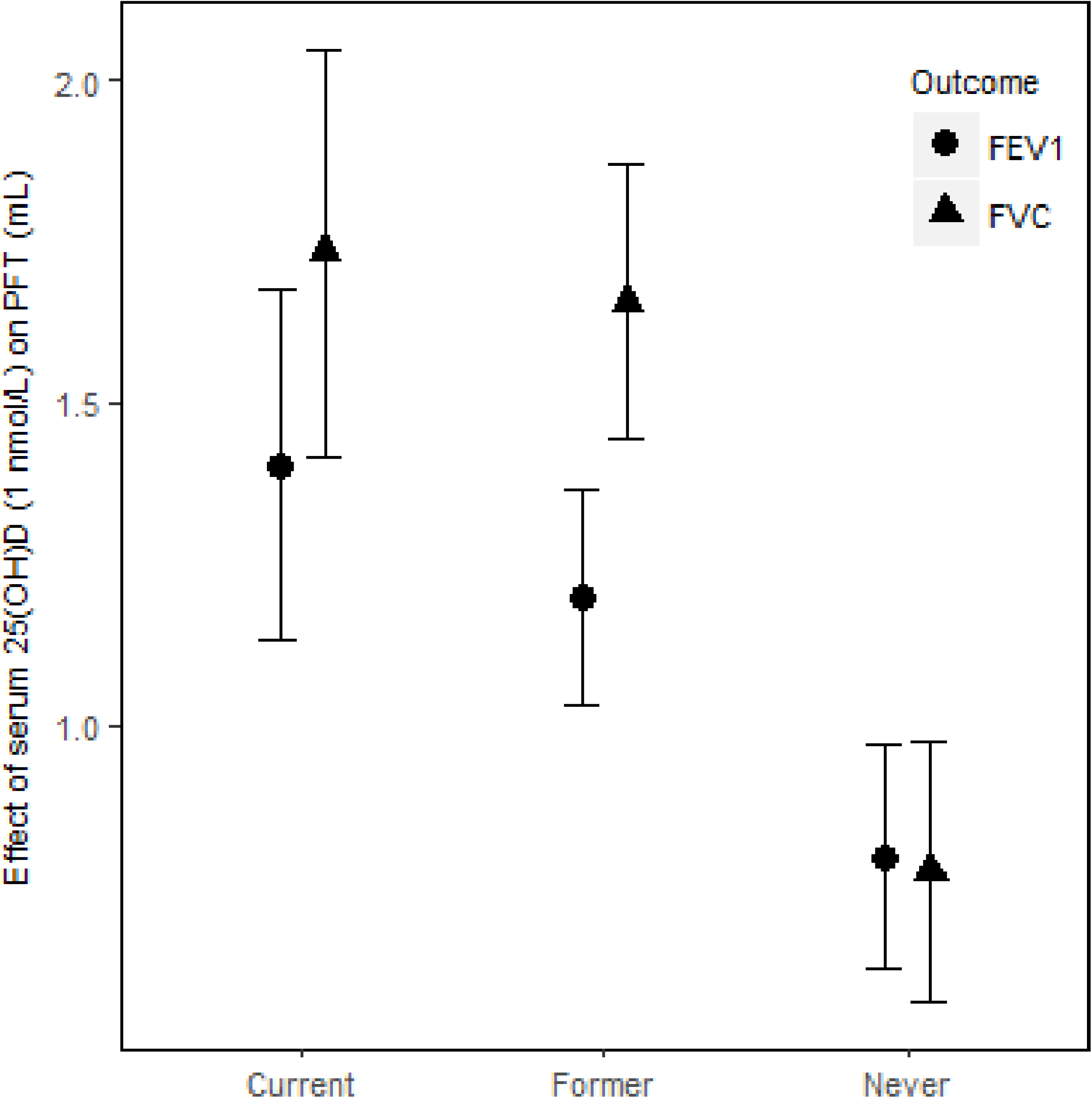
Meta-analysis of the association of serum 25(OH)D–PFT outcomes among current, former, and never smokers in the European ancestry cohorts in the CHARGE Consortium. FEV_1_ and FVC are presented for each smoking status. β (unit: mL) denotes that 1 nmol/L higher serum 25(OH)D was associated with a β mL higher FEV_1_ (or FVC), calculated from an analysis including the interaction of serum 25(OH)D and smoking status. The error bar represents ± 1 standard error. We used 22,787 EA participants for the FEV_1_ outcome and 22,777 EA participants for the FVC outcome. Abbreviation: 25(OH)D, 25-Hydroxyvitamin D; EA, European Ancestry; FEV_1_, Forced Expiratory Volume in the First Second; FVC, Forced Vital Capacity; PFT, Pulmonary Function Test.

To examine the potential impact of family relatedness between the FHS-Gen3 and the FHS-Offspring cohorts on the meta-analysis, sensitivity analysis confirmed that the findings were unchanged when either cohort was excluded (results not shown).

Cohort-specific findings (**Supplemental Table 5 and 6**) from models that included the 25(OH)D × smoking status interaction terms were combined in secondary meta-analyses (**Supplemental Table 7**). In the EA cohorts, 25(OH)D had a greater positive association with FVC in current smokers than in never smokers (β_current × 25(OH)D_ = 7.5 mL for 10 nmol/L increment of 25(OH)D, P = 0.047). Similarly, 25(OH)D had a greater positive association with FVC in former smokers than in never smokers (β_former × 25(OH)D_ = 7.9 mL for 10 nmol/L increment of 25(OH)D, P = 0.0065) (**Figure 3**). For the FEV_1_ outcome in the EA cohorts, the interaction coefficients for 25(OH)D and smoking status had the same positive direction as the coefficients for FVC, but were not statistically significant for either current (P = 0.14) or former smokers (P = 0.14). There was no statistical evidence of interaction of 25(OH)D and cigarette smoking in the AA cohorts for either outcome. To put the interaction finding into context, a 10 nmol/L higher serum 25(OH)D was associated with a 17.3 mL higher FVC in current smokers and a 16.6 mL higher FVC in former smokers, which was more than double the association magnitude in never smokers (β = 7.8 mL). A similar trend was found for the FEV_1_ outcome in the EA cohorts. For 10 nmol/L higher serum 25(OH)D, FEV_1_ was higher by 14.0 mL in current smokers, 12.0 mL in former smokers, and 8.0 mL in never smokers (**Figure 4**).

## DISCUSSION

This study investigated the association of serum 25(OH)D with pulmonary function using multiple cohorts of different ancestries. We found a consistently positive association of serum 25(OH)D with FEV_1_ and FVC across both EA and AA groups. In addition, in the EA group, a significantly stronger association was observed for current and former smokers, compared to never smokers.

A previous cross-sectional study in a European ancestry population (two Copenhagen cohorts: *n* = 10,116 and *n* = 8,391 respectively) similarly reported positive associations of 25(OH)D with FEV_1_ percentage predicted and FVC percentage predicted, but not with FEV_1_/FVC^(16)^. The magnitude of the association was about four times greater in the Copenhagen study, which may be due to the difference in the mean serum 25(OH)D (Danish median ∼42 nmol/L vs. CHARGE median of ∼65 nmol/L) given our finding that the 25(OH)D–PFT association was stronger in cohorts with lower serum 25(OH)D. Our finding for the serum 25(OH)D–FEV_1_ association was similar in magnitude to the association reported in a British cohort of 6,789 participants with an average age of 45 years^(17)^, but weaker than a previous report from the FHS cohort^(15)^. Given that the rate of decline in FEV_1_ at age 45 is increased by ∼15 mL/year in current smokers^(31)^, we estimate that a 10 nmol/L higher 25(OH)D is similar to approximately 1 year of current smoking-related decline in FEV_1_ for both ancestries, but in the opposite direction. Potential biological mechanisms for a causal association between low 25(OH)D levels and low pulmonary function include an altered immune response that increases susceptibility to inflammation, a reduction in pulmonary parenchyma related to extracellular matrix homeostasis important for lung structure, and/or a decrease in serum calcium that could adversely affect thoracic skeleton mobility and respiratory muscle performance^(32)^.

Our findings show that the association of serum 25(OH)D with FEV_1_ and FVC were stronger in magnitude in AA versus EA participants, although the difference by race did not reach statistical significance. The finding may reflect the lower serum 25(OH)D in AA participants, which is consistent with the meta-regression finding and with a previous study reporting attenuated associations at higher serum 25(OH)D (15). Future studies that investigate genetic variation in EAs and AAs in the context of serum 25(OH)D may help explain the differences.

In EA participants, the positive interaction terms between serum 25(OH)D and smoking status supported a stronger magnitude of association of serum 25(OH)D with FVC in current and former smokers than in never smokers, with a consistent, but not statistically significant, difference for FEV_1_. The interaction finding is consistent with a prior cross-sectional National Health and Nutrition Examination Survey (NHANES) study, which reported a stronger 25(OH)D–FEV_1_ association in current and former smokers than in never smokers that was near statistical significance (P = 0.06)^(13)^. Given smokers have a higher level of oxidative stress and lower pulmonary function than never smokers, partly due to chronic inflammation in lung tissue, the stronger protective association of 25(OH)D on pulmonary function in smokers suggests a benefit for smokers. To explore this interaction, estimates of the 25(OH)D–PFT association were computed within each smoking category. In EA participants, the 25(OH)D–FEV_1_ (or FVC) associations were statistically significant in all strata. Generally, in ever smokers of European ancestry, the coefficients for 25(OH)D were greater for FVC than for FEV_1_.

Meta-regression provided additional evidence for effect modification by smoking. The proportion of ever smokers was a significant modifier of the association of serum 25(OH)D with FEV_1_ and FVC. The higher the proportion of ever smokers, the greater the 25(OH)D–PFT association. More specifically, the proportion of former smokers explained the heterogeneity in the 25(OH)D–PFT association across cohorts more fully than the proportion of current smokers; this may be explained by a survival bias in older participants who were current smokers. The meta-regression based on mean age of the cohorts yielded findings that were consistent with a prior NHANES study that showed the association of 25(OH)D and FEV_1_ was greater in people over age 60 compared to younger individuals^(13)^.

There are several methodological considerations in interpreting the findings of this study. First, the meta-regression showed stronger 25(OH)D–PFT associations in cohorts with lower mean serum 25(OH)D, indicating a non-linear 25(OH)D–PFT association. This finding is consistent with a prior study in the FHS cohort, which reported a non-linear association and a stronger 25(OH)D–FEV_1_ association in participants at risk of vitamin D deficiency (< 30 nmol/L)^(15)^. Second, serum 25(OH)D was measured by four different methods across the cohorts. For example, two cohorts with high mean 25(OH)D (> 90 nmol/L) used RIA methods. These same cohorts had a lower magnitude estimate of the 25(OH)D–PFT association; if the higher mean represents the ‘truth’ (and is not caused by measurement error in the RIA assay), then the lower 25(OH)D–PFT association may be primarily driven by the vitamin D distribution and not by the RIA method. Whether the assay method itself directly influences the estimate of the 25(OH)D– PFT association requires further data. Third, in this cross-sectional meta-analysis, there were minor differences in the time separation between the measurement of serum 25(OH)D and pulmonary function, but the meta-regression test for heterogeneity confirmed that time separation between measurements did not affect the 25(OH)D–PFT associations. Indeed, past studies with longitudinal measurements of serum 25(OH)D reported a high correlation of 25(OH)D measurements over a long period of time, with a correlation coefficient of 0.7 for measurements separated by 1 year, 0.5 for measurements separated by 5 years^(33)^, and 0.42-0.52 for measurements separated by 14 years^(34)^, which supports the use of a single 25(OH)D measurement to represent usual level. Fourth, residual confounding was unlikely given the consistent results across multiple cohorts in various settings. Weight was adjusted for the FVC outcome, given that higher weight and adiposity negatively affects lung volume (i.e., FVC)^(35)^; weight was not adjusted in the FEV_1_ models, given FEV_1_ is a measure of airways obstruction and not physical restriction of lung volume. Physical activity was not adjusted because it is not a confounder in estimating the serum 25(OH)D–PFT association; while physical activity is known to contribute to oxygen utilization in lungs^(36)^ there is little evidence and no biological rationale for a causal association of physical activity with either FEV_1_ or FVC^(37)^, which are markers for airways obstruction and lung volume, respectively. Finally, we do not expect selection bias to affect the estimate of the serum vitamin D–PFT association in this meta-analysis; the association magnitude and direction was consistent across all cohorts, regardless of the proportion of the original cohort contributing to the analysis. Thus, selection bias is expected to be negligible and would likely lead to an underestimated association, given the participants retained in the cohorts are expected to be, on average, healthier than those who were lost to follow-up.

This study meta-analyzed the serum 25(OH)D–PFT association across nine cohorts, according to a common pipeline that harmonized the variables and statistical analysis. The sample size comprised 17,569 EA participants from the United States; 5,269 EA participants from Iceland and the Netherlands; and 4,290 AA participants from the United States, all of whom were 19 to 95 years old. The sample provided excellent representation of the U.S. population, based on comparisons of demographic factors including sex, height, weight, smoking status, and COPD prevalence (∼6.1%) to national surveys^(38^-^40)^, which strengthens the external validity of the study’s findings.

In summary, using meta-analysis, we estimated a positive association of serum 25(OH)D with the pulmonary function parameters FEV_1_ and FVC in both EA and AA participants. Associations varied by smoking status in the EA group, with stronger serum 25(OH)D–PFT associations seen in current and former smokers. The observational design means we cannot infer a causal association, and future studies, such as randomized controlled trials or Mendelian randomization studies, are needed to further investigate the causality of 25(OH)D on pulmonary function.

## ACKNOWLEDGMENTS

We thank Prof. James Booth in the Department of Biological Statistics and Computational Biology at Cornell University for providing advice on meta-analysis. We thank the staff (Lynn Johnson and Francoise Vermeylen) in Cornell Statistical Consulting Unit for their role in providing advice on methods.

## CONTRIBUTORS

PAC, DBH, and JX conceived and designed the study. RGB, JL, JD, SAG, LL, SJL, KEN, AVS, BMP, and LMS provided the data and supervised the data analysis in each cohort. JX, TMB, RRR, AVS, AWM, FS, NT, and XZ analyzed data within each cohort. JX, PAC and DBH meta-analyzed and interpreted the data. JX, PAC and DBH co-wrote and edited the first draft of the manuscript. PAC, DBH and JX had primary responsibility for final content. All authors provided data, analytic support and/or study design suggestions at all stages, critically reviewed the manuscript, and read and approved the final version.

## REFERENCES

1. Minino AM, Murphy SL, Xu J, et al. (2011) Deaths: final data for 2008. Natl Vital Stat Rep 59, 1–126.

2. Collaborators GBDCoD (2017) Global, regional, and national age-sex specific mortality for 264 causes of death, 1980-2016: a systematic analysis for the Global Burden of Disease Study 2016. Lancet 390, 1151–210.

3. Weiss ST (2010) Lung function and airway diseases. Nat Genet 42, 14–6.

4. Forey BA, Thornton AJ, Lee PN (2011) Systematic review with meta-analysis of the epidemiological evidence relating smoking to COPD, chronic bronchitis and emphysema. BMC Pulm Med 11, 36.

5. Guerra S, Stern DA, Zhou M, et al. (2013) Combined effects of parental and active smoking on early lung function deficits: a prospective study from birth to age 26 years. Thorax 68, 1021–8.

6. Fletcher JM, Basdeo SA, Allen AC, et al. (2012) Therapeutic use of vitamin D and its analogues in autoimmunity. Recent Pat Inflamm Allergy Drug Discov 6, 22–34.

7. Litonjua AA, ebrary Inc. (2012) Vitamin D and the lung mechanisms and disease associations. New York: Humana Press/Springer, http://proxy.library.cornell.edu/login?url=http://link.springer.com/openurl?genre=book&isbn=978-1-61779-887-0.

8. Bikle DD, Gee E, Halloran B, et al. (1986) Assessment of the free fraction of 25-hydroxyvitamin D in serum and its regulation by albumin and the vitamin D-binding protein. J Clin Endocrinol Metab 63, 954–9.

9. Nykjaer A, Dragun D, Walther D, et al. (1999) An endocytic pathway essential for renal uptake and activation of the steroid 25-(OH) vitamin D3. Cell 96, 507–15.

10. Zerwekh JE (2008) Blood biomarkers of vitamin D status. Am J Clin Nutr 87, 1087S–91S.

11. Nielson CM, Jones KS, Chun RF, et al. (2016) Free 25-Hydroxyvitamin D: Impact of Vitamin D Binding Protein Assays on Racial-Genotypic Associations. J Clin Endocrinol Metab 101, 2226–34.

12. Freedman BI, Register TC (2012) Effect of race and genetics on vitamin D metabolism, bone and vascular health. Nat Rev Nephrol 8, 459–66.

13. Black PN, Scragg R (2005) Relationship between serum 25-hydroxyvitamin d and pulmonary function in the third national health and nutrition examination survey. Chest 128, 3792–8.

14. Choi CJ, Seo M, Choi WS, et al. (2013) Relationship between serum 25-hydroxyvitamin D and lung function among Korean adults in Korea National Health and Nutrition Examination Survey (KNHANES), 2008-2010. J Clin Endocrinol Metab 98, 1703–10.

15. Hansen JG, Gao W, Dupuis J, et al. (2015) Association of 25-Hydroxyvitamin D status and genetic variation in the vitamin D metabolic pathway with FEV1 in the Framingham Heart Study. Respir Res 16, 81.

16. Afzal S, Lange P, Bojesen SE, et al. (2014) Plasma 25-hydroxyvitamin D, lung function and risk of chronic obstructive pulmonary disease. Thorax 69, 24–31.

17. Berry DJ, Hesketh K, Power C, et al. (2011) Vitamin D status has a linear association with seasonal infections and lung function in British adults. Br J Nutr 106, 1433–40.

18. Thuesen BH, Skaaby T, Husemoen LL, et al. (2015) The association of serum 25-OH vitamin D with atopy, asthma, and lung function in a prospective study of Danish adults. Clin Exp Allergy 45, 265–72.

19. Tolppanen AM, Williams D, Henderson J, et al. (2011) Serum 25-hydroxy-vitamin D and ionised calcium in relation to lung function and allergen skin tests. Eur J Clin Nutr 65, 493–500.

20. Shaheen SO, Jameson KA, Robinson SM, et al. (2011) Relationship of vitamin D status to adult lung function and COPD. Thorax 66, 692–8.

21. Niruban SJ, Alagiakrishnan K, Beach J, et al. (2015) Association between vitamin D and respiratory outcomes in Canadian adolescents and adults. J Asthma 52, 653–61.

22. Tolppanen AM, Sayers A, Granell R, et al. (2013) Prospective association of 25-hydroxyvitamin d3 and d2 with childhood lung function, asthma, wheezing, and flexural dermatitis. Epidemiology 24, 310–9.

23. Lange NE, Sparrow D, Vokonas P, et al. (2012) Vitamin D deficiency, smoking, and lung function in the Normative Aging Study. Am J Respir Crit Care Med 186, 616–21.

24. Khan S, Mai XM, Chen Y (2013) Plasma 25-hydroxyvitamin D associated with pulmonary function in Canadian adults with excess adiposity. Am J Clin Nutr 98, 174–9.

25. O’Neal JD (2015) Vitamin D supplementation regimens for HIV-infected patients: a historical chart review. Scholar Archive.

26. Mannino DM, Diaz-Guzman E, Buist S (2011) Pre-and post-bronchodilator lung function as predictors of mortality in the Lung Health Study. Respir Res 12, 136.

27. Phinney KW (2008) Development of a standard reference material for vitamin D in serum. Am J Clin Nutr 88, 511S–2S.

28. Fujiyoshi A, Polgreen LE, Hurley DL, et al. (2013) A cross-sectional association between bone mineral density and parathyroid hormone and other biomarkers in community-dwelling young adults: the CARDIA study. J Clin Endocrinol Metab 98, 4038–46.

29. Higgins JP, Thompson SG, Deeks JJ, et al. (2003) Measuring inconsistency in meta-analyses. BMJ 327, 557–60.

30. Borenstein M, Wiley InterScience (Online service). (2009) Introduction to meta-analysis. Chichester, U.K.: John Wiley & Sons, http://encompass.library.cornell.edu/cgi-bin/checkIP.cgi?access=gateway_standard%26url=http://onlinelibrary.wiley.com/book/10.1002/9780470743386.

31. Mirabelli MC, Preisser JS, Loehr LR, et al. (2016) Lung function decline over 25 years of follow-up among black and white adults in the ARIC study cohort. Respir Med 113, 57–64.

32. Herr C, Greulich T, Koczulla RA, et al. (2011) The role of vitamin D in pulmonary disease: COPD, asthma, infection, and cancer. Respir Res 12, 31.

33. Hofmann JN, Yu K, Horst RL, et al. (2010) Long-term variation in serum 25-hydroxyvitamin D concentration among participants in the Prostate, Lung, Colorectal, and Ovarian Cancer Screening Trial. Cancer Epidemiol Biomarkers Prev 19, 927–31.

34. Jorde R, Sneve M, Hutchinson M, et al. (2010) Tracking of serum 25-hydroxyvitamin D levels during 14 years in a population-based study and during 12 months in an intervention study. Am J Epidemiol 171, 903–8.

35. Amara CE, Koval JJ, Paterson DH, et al. (2001) Lung function in older humans: the contribution of body composition, physical activity and smoking. Ann Hum Biol 28, 522–36.

36. Burri PH, Gehr P, Muller K, et al. (1976) Adaptation of the growing lung to increased VO2. I. IDPN as inducer of hyperactivity. Respir Physiol 28, 129–40.

37. Cheng YJ, Macera CA, Addy CL, et al. (2003) Effects of physical activity on exercise tests and respiratory function. Br J Sports Med 37, 521–8.

38. Jamal A, Homa DM, O’Connor E, et al. (2015) Current cigarette smoking among adults - United States, 2005-2014. MMWR Morb Mortal Wkly Rep 64, 1233–40.

39. Fryar CD, Gu Q, Ogden CL (2012) Anthropometric reference data for children and adults: United States, 2007-2010. Vital Health Stat 11, 1–48.

40. Ward BW, Nugent CN, Blumberg SJ, et al. (2017) Measuring the Prevalence of Diagnosed Chronic Obstructive Pulmonary Disease in the United States Using Data From the 2012-2014 National Health Interview Survey. Public Health Rep 132, 149–56.

